## Supplementary material for "De novo identification of universal cell mechanics gene signatures": Figure Supplements

Figure Supplements for

BioRχiv 2021.04.26.441418 V4

This file includes the following figure supplements:

**Figure 1–figure supplement 1** – Characterization of mechanical cell properties using real-time deformability cytometry (RT-DC).

**Figure 2–figure supplement 1** – Plots of area vs deformation for different cell states in the characterized systems.

**Figure 3–figure supplement 1** – Gene ontology (GO) enrichment analysis of obtained target genes.

**Figure 4–figure supplement 1** – Expression of identified target genes in the CCLE microarray dataset used for validation.

**Figure 4–figure supplement 2** – Expression of identified target genes in the CCLE RNA-Seq dataset used for validation.

**Figure 4–figure supplement 3** – Expression of identified target genes in the Genentech dataset used for validation.

**Figure 4–figure supplement 4** – Relation between the magnitude of apparent Young's modulus change and the absolute change in the expression levels of target genes.

**Figure 4–figure supplement 5** – ROC curves characterizing classification performance of the five genes from the conserved module.

**Figure 5–figure supplement 1** – CAV1 knock-out mouse embryonic fibroblasts (CAV1KO) have lower stiffness compared to the wild type cells (WT).

**Figure 5–figure supplement 2** – Apparent Young's modulus values across the probing frequencies characteristic for the three measurement methods.

**Figure 5–figure supplement 3** – Plots of area vs deformation from RT-DC measurements of cells with perturbed CAV1 levels.

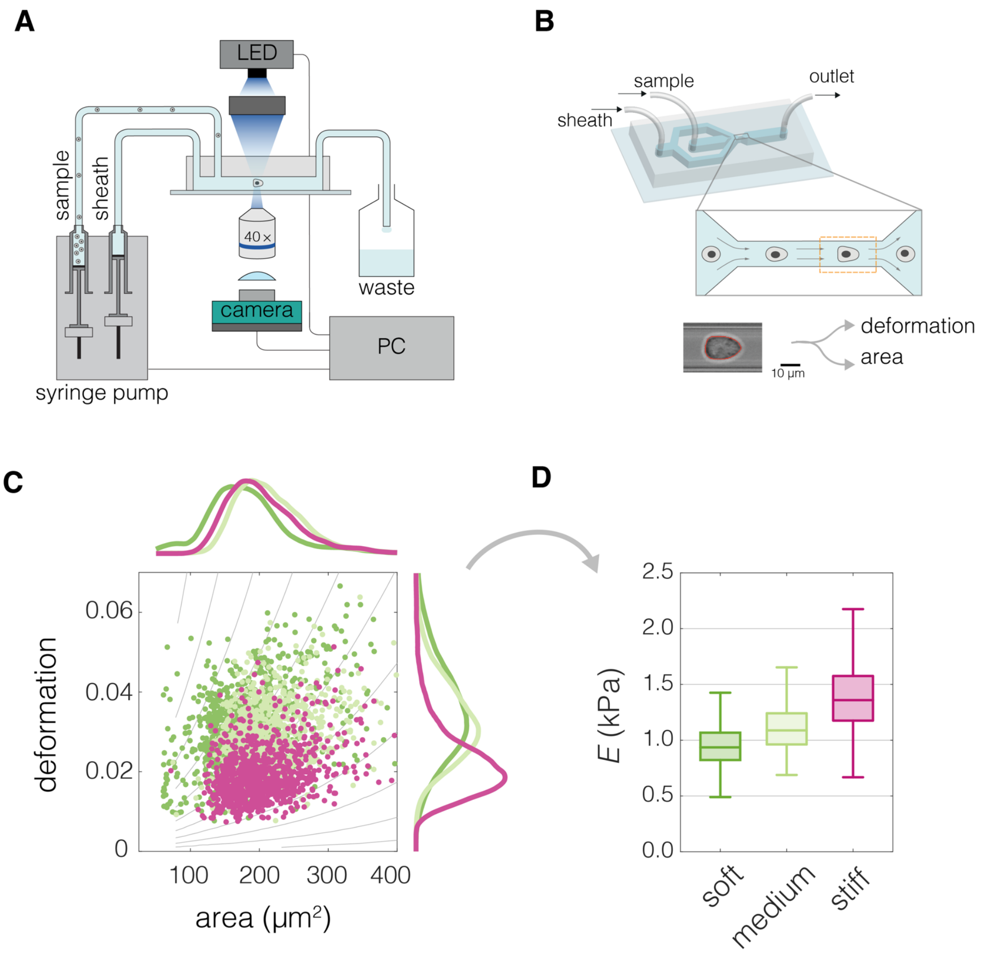


#### Figure 1–figure supplement 1

**Characterization of mechanical cell properties using real-time deformability cytometry (RT-DC).**

(**A**) Schematic overview of the RT-DC setup. Computer-operated syringe pumps flow the cell-containing sample as well as the sheath fluid into the microfluidic chip. Imaging of the cells deformed in the microfluidic channel is performed at 2,000 frames per second using an LED-based stroboscopic illumination and a CMOS camera. (**B**) 3D illustration of the microfluidic chip used for the RT-DC measurements, close-up depicts the constriction of the channel in which cells are deformed, the imaged region of interest is indicated by an orange dashed line. At the bottom an exemplary image of a cell is shown. A contour is fitted to the cell in real time (marked in red), based on which cell area and deformation are calculated. (**C**) An exemplary plot of deformation versus area of three different cell populations. The gray isoelasticity lines in the background indicate regions of the same apparent Young's moduli. (**D**) Box plot of apparent Young's modulus, *E*, estimated based on deformation and area in (C). The cell population with same area but higher deformation has lower *E* (bright green compared to magenta). For cells with similar deformation, the one of smaller area has lower *E* (dark green compared to bright green). The exemplary data in (C) and (D) corresponds to exemplary measurements of Wa-hT (dark green), EBC1 (bright green) and A549 (magenta) cell lines. The box plots in (D) spread from 25th to 75th percentiles with a line at the median, whiskers span 1.5 × interquartile range (IQR).

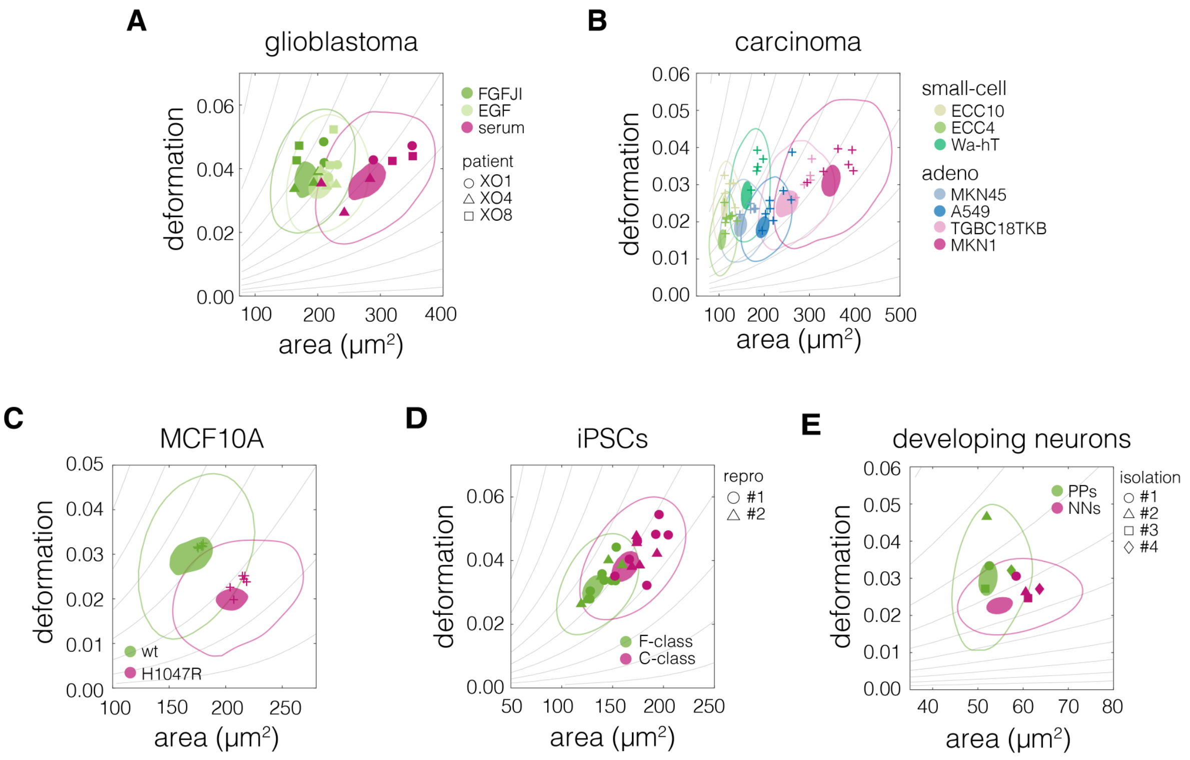


#### Figure 2–figure supplement 1

**Plots of area vs deformation for different cell states in the characterized systems.**

Panels correspond to the following systems: (**A**) glioblastoma, (**B**) carcinoma, (**C**) non-tumorigenic breast epithelia MCF10A, (**D**) induced pluripotent stem cells (iPSCs), and (**E**) developing neurons. 95%- and 50% density contours of data pooled from all measurements of given cell state are indicated by shaded areas and continuous lines, respectively. Datapoints indicate medians of individual measurements. The symbol shapes represent cell lines derived from three different patients (A), two different reprogramming series (D), and three different cell isolations (E), as indicated in the respective panels. The isoelasticity lines in the background (gray) indicate regions of the same apparent Young's moduli. PPs – proliferating progenitors, NNs – newborn neurons.

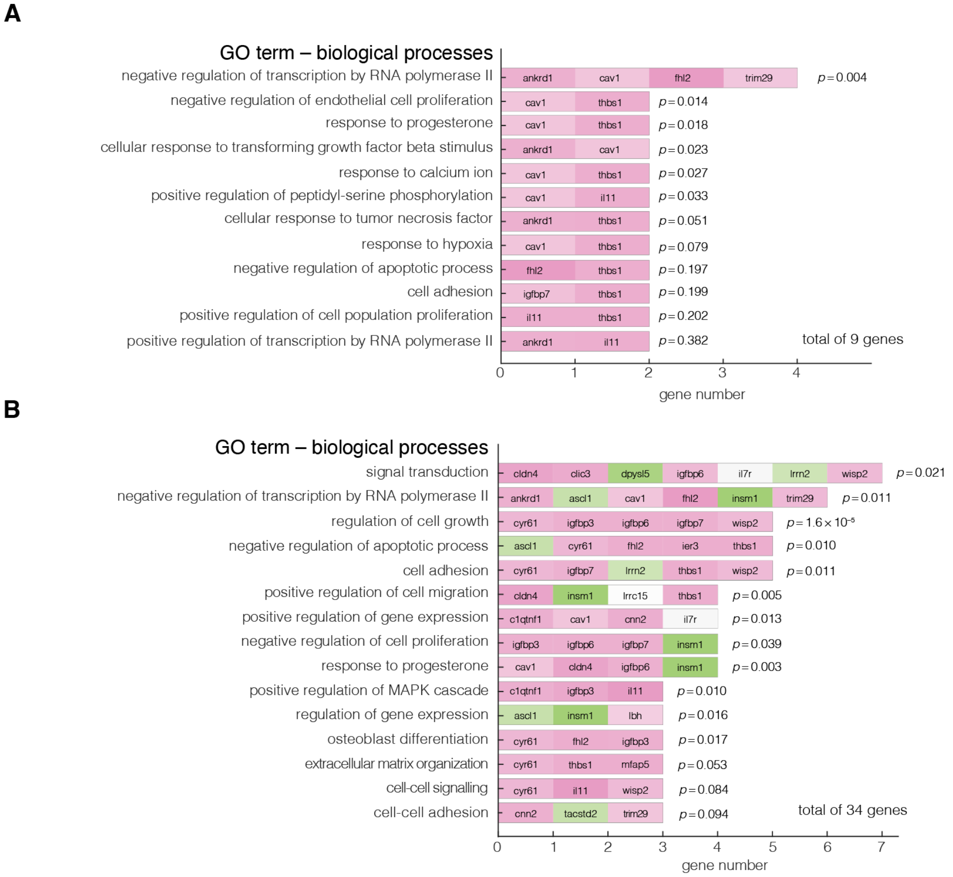


### Figure 3–figure supplement 1

**Gene ontology (GO) enrichment analysis of obtained target genes.**

Enriched GO terms of biological processes are summarized for: (**A**) 9 genes corresponding to the results from **Figure 3G**, (**B**) 34 genes corresponding to all nodes presented in **Figure 3, E–G**. The analysis was performed using *DAVID 6.8* functional annotation tool online, with *Homo sapiens* as background dataset, ENSMBL gene IDs as input, and focused on direct GO terms for biological processes. Color code of the blocks corresponds to the level of expression in stiff states with green corresponding to low expression and magenta corresponding to high expression. The reported *p* values are the Fisher's exact *p* values obtained using a two tailed two sample *t*-test.

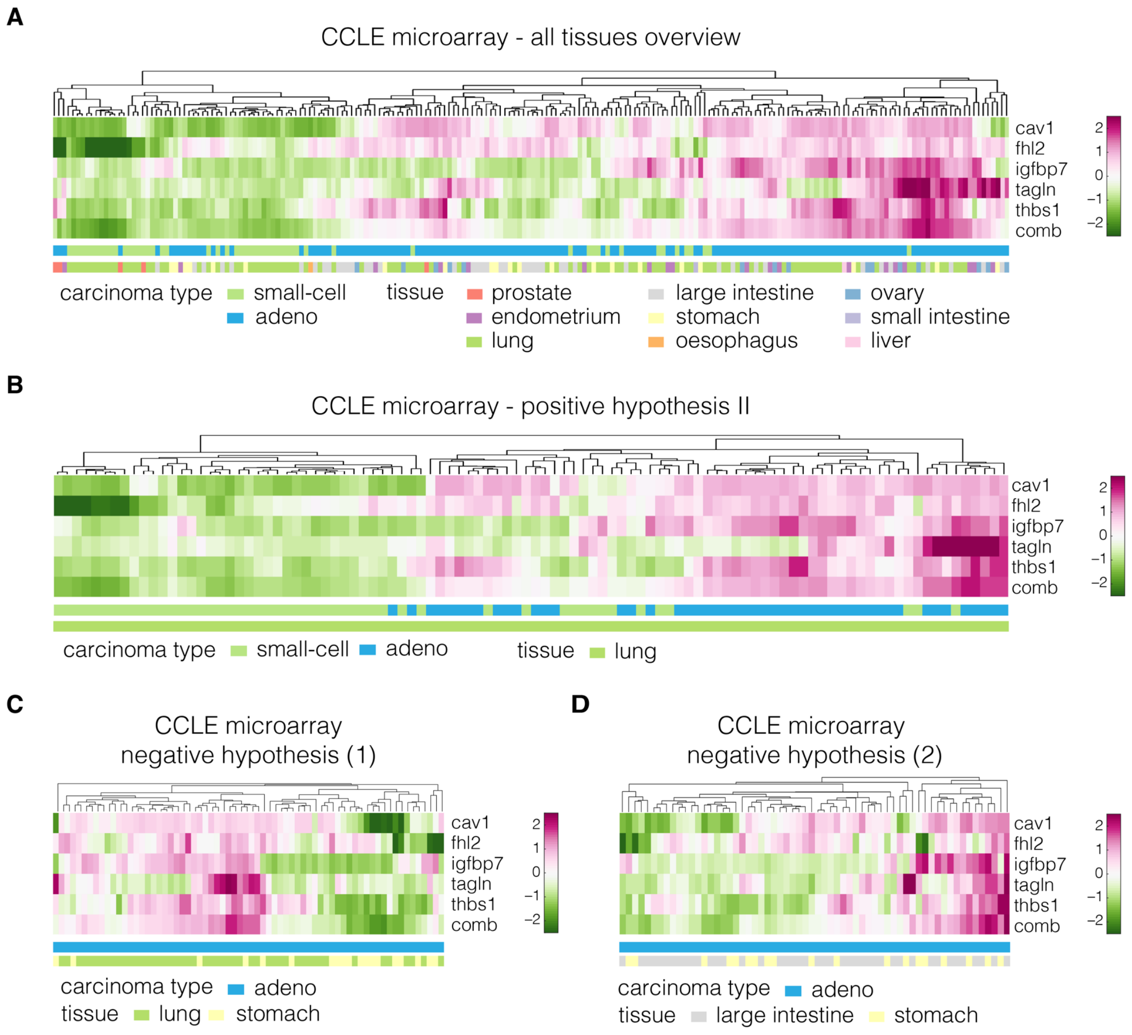


#### Figure 4–figure supplement 1

**Expression of identified target genes in the CCLE microarray dataset used for validation.**

Panels show unsupervised clustering heat maps of expression data from the CCLE microarray dataset and include: (**A**) adeno and small-cell carcinoma samples across all tissues present in the dataset, (**B**) adeno and small-cell carcinoma samples corresponding to lung tissue (used for testing of the positive hypothesis II, see **Table 3** in the main text), (**C**) adenocarcinoma samples corresponding to lung and stomach (used for testing of negative hypothesis, see **Table 3** in the main text), (**D**) adenocarcinoma samples corresponding to large intestine and stomach (used for testing of the negative hypothesis, see **Table 3** in the main text). Clustering was performed using clustergram function in MATLAB (R2020a, MathWorks) on log-normalized expression data. The bars under each heatmap are color-coded for the carcinoma type and tissue of origin (top and bottom bar, respectively) as specified in panel legends. Sample IDs corresponding to each class are listed in the **Supplementary File 3.**

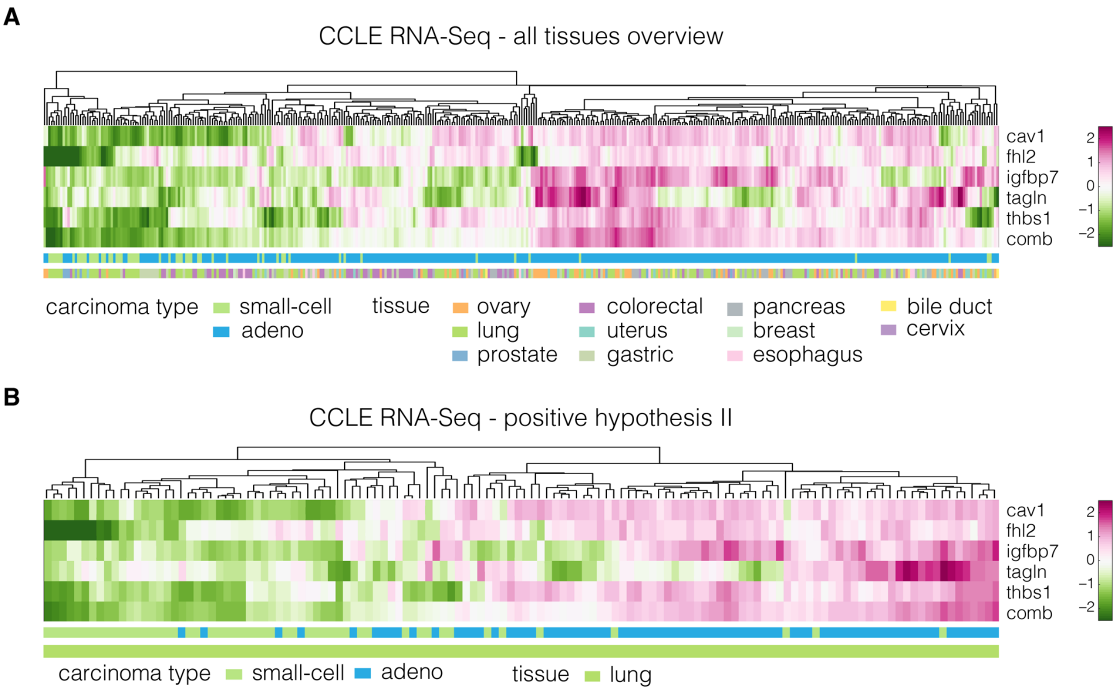


#### Figure 4–figure supplement 2

**Expression of identified target genes in the CCLE RNA-Seq dataset used for validation.**

Panels show unsupervised clustering heat maps of expression data from the CCLE RNA-Seq dataset and include: (**A**) adeno and small-cell carcinoma samples across all tissues present in the dataset, (**B**) adeno and small-cell carcinoma samples corresponding to lung tissue (used for testing of the positive hypothesis II, see **Table 3** in the main text). Clustering was performed using clustergram function in MATLAB (R2020a, MathWorks) on log-normalized expression data. The bars under each heatmap are color-coded for the carcinoma type and tissue of origin (top and bottom bar, respectively) as specified in panel legends. Sample IDs corresponding to each class are listed in the **Supplementary File 3**.

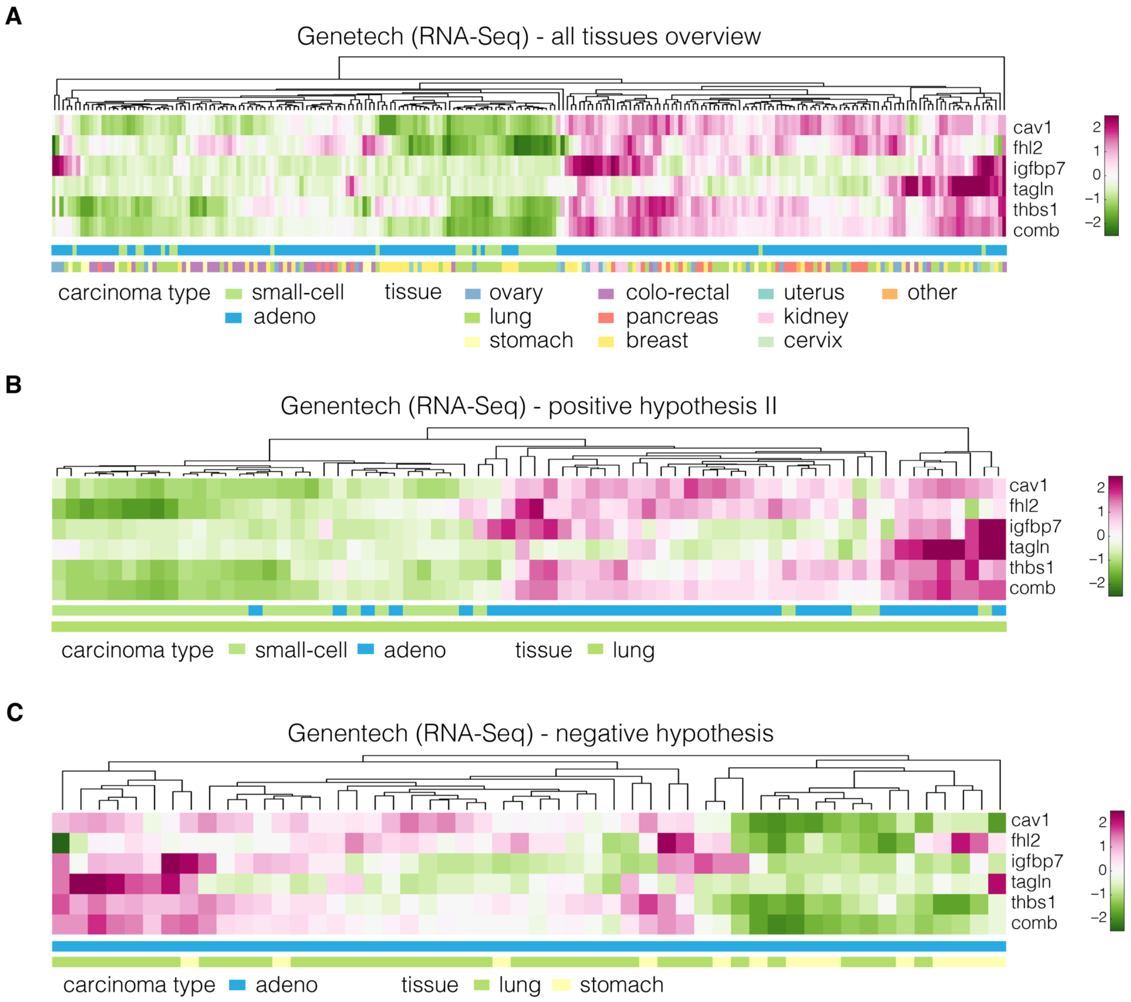


#### Figure 4–figure supplement 3

**Expression of identified target genes in the Genentech dataset used for validation.**

Panels show unsupervised clustering heat maps of expression data from the Genentech dataset and include: (**A**) adeno and small-cell carcinoma samples across all tissues present in the dataset, (**B**) adeno and small-cell carcinoma samples corresponding to lung tissue (used for testing of the positive hypothesis II, see **Table 3** in the main text), (**C**) adenocarcinoma samples corresponding to lung and stomach (used for testing of the negative hypothesis, see **Table 3** in the main text). Clustering was performed using clustergram function in MATLAB (R2020a, MathWorks) on log-normalized expression data. The bars under each heatmap are color-coded for the carcinoma type and tissue of origin (top and bottom bar, respectively) as specified in panel legends. Sample IDs corresponding to each class are listed in the **Supplementary File 3.**

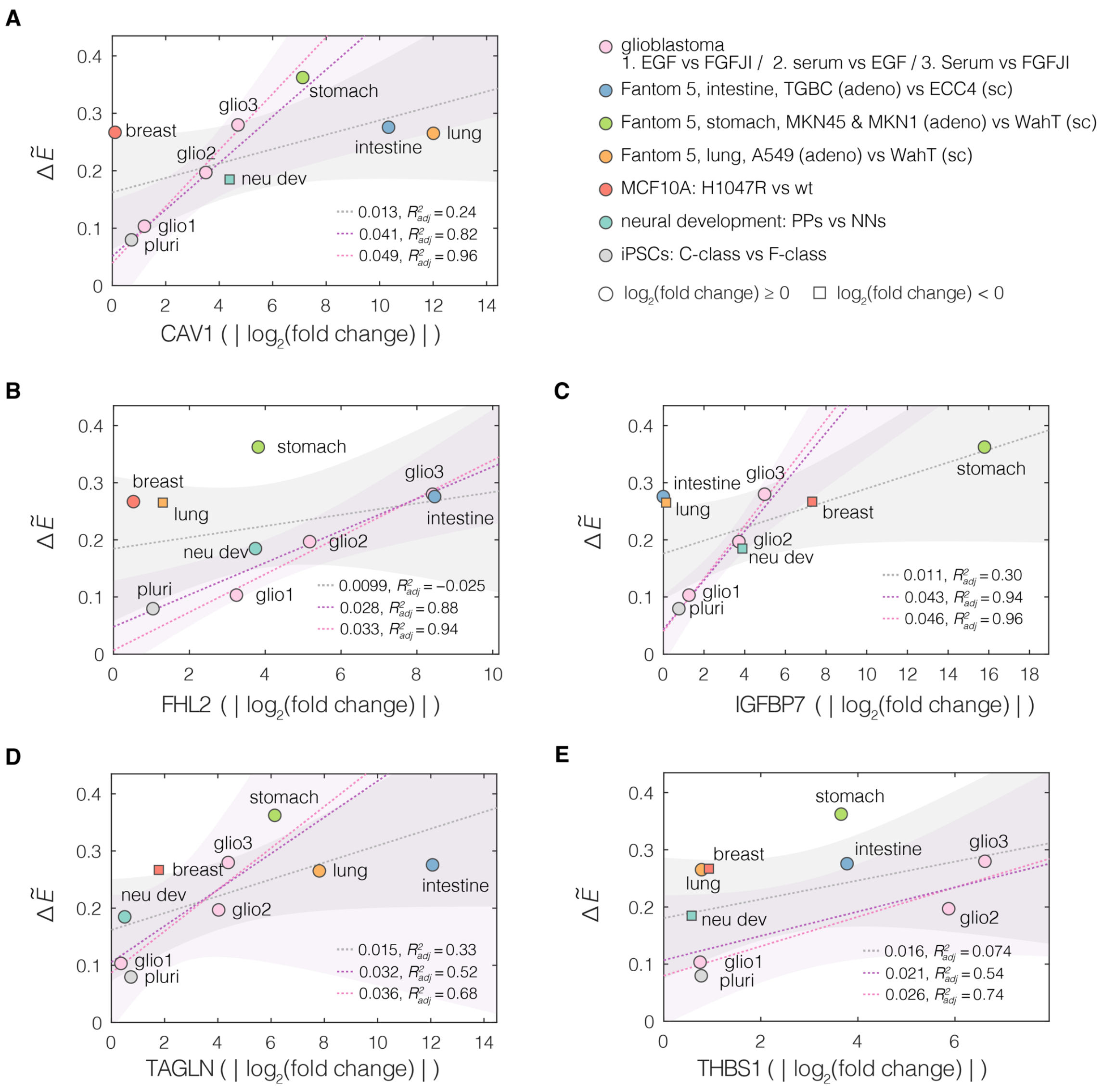


#### Figure 4–figure supplement 4

**Relation between the magnitude of apparent Young's modulus change and the absolute change in the expression levels of target genes.**

Plots of normalized change in apparent Young's modulus $\Delta\tilde{E}$ versus absolute value of change in expression for the target genes from conserved module: (**A**) CAV1, (**B**) FHL2, (**C**) IGFBP7, (**D**) TAGLN, and (**E**) THBS1. Every soft-stiff state pair from the respective datasets is presented as an individual point. $\Delta\tilde{E} = \frac{E_{stiff}-E_{soft}}{E_{stiff}}$, where $E_{stiff}$ and $E_{soft}$ correspond to the apparent Young's moduli (mean of all measurements) of the stiff and soft states within the given pairs, respectively. The dotted lines correspond to linear fits to all (gray), similar lineage (glioblastoma, developing neurons and iPSC; purple) and glioblastoma only (pink) datapoints. Shaded areas represent 95% confidence intervals (displayed for all and similar lineage fits only). The slope of the respective linear fits and the adjusted $R^2$, $R^2_{adj}$, are reported in the plots. For the linage-selected data, the fitted slopes become higher and the quality of the fits better (higher $R^2_{adj}$).

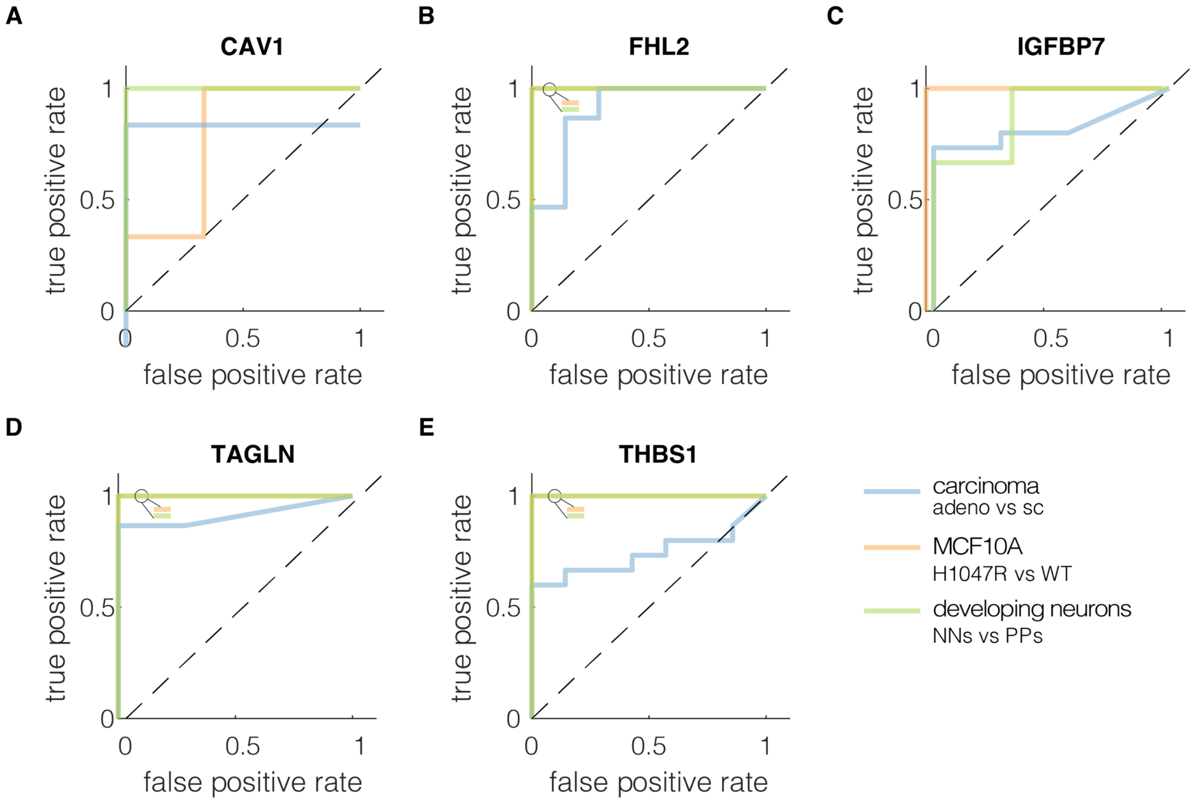


#### Figure 4—figure supplement 5

**ROC curves characterizing classification performance of the five genes from the conserved module.**

True positive rate was plotted against the false positive rate at different classification thresholds for each soft-stiff phenotype pair from the validation datasets for: (**A**) CAV1, (**B**) FHL2, (**C**) IGFBP7, (**D**) TAGLN, and (**E**) THBS1. The insets in the upper left corners of the plot show the colors of all overlying curves with AUC = 1. The ROC curves were constructed using *perfcurve* function in *MATLAB* (*R2020a,* MathWorks). adeno – adenocarcinoma, sc – small cell carcinoma, WT – wild type, PPs – proliferating progenitors, NNs – newborn neurons.

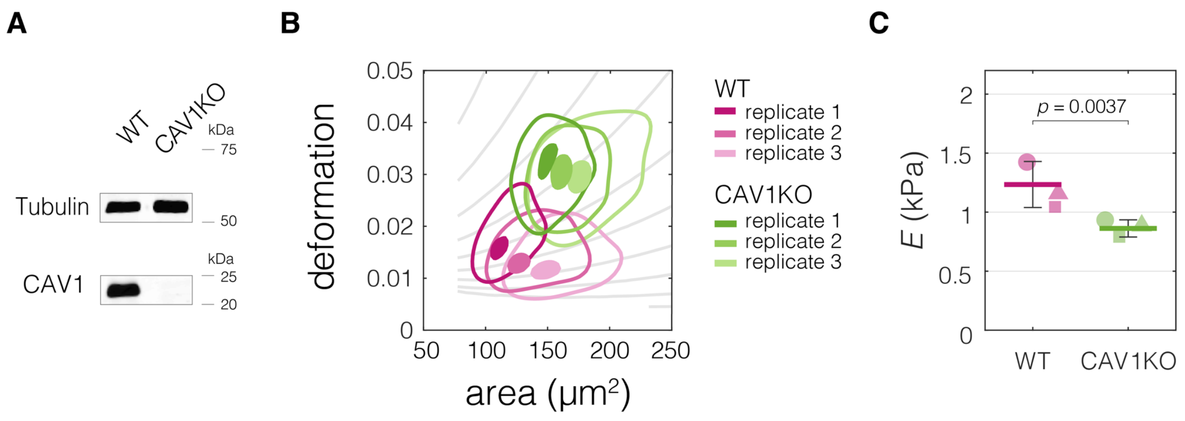


#### Figure 5–figure supplement 1

**CAV1 knock-out mouse embryonic fibroblasts (CAV1KO) have lower stiffness compared to the wild type cells (WT).**

(**A**) Western blot analysis of CAV1 expression levels in CAV1KO compared to WT cells. (**B**) Plots of area vs deformation for CAV1KO and WT cells characterized with RT-DC. Contour plots delineate 95% and 50% density areas (solid lines and filled area, respectively) of data from individual measurement replicates ($n$ = 3). The isoelasticity lines in the background (gray) indicate regions of the same apparent Young's moduli. (**C**) Apparent Young's modulus values estimated for WT and CAV1KO cells using area-deformation data in (B). The Symbol shapes represent experimental replicates. Horizontal lines delineate medians with mean absolute deviation (MAD) as error, datapoints represent medians of the individual replicates. Statistical analysis was performed using generalized linear mixed effects model.

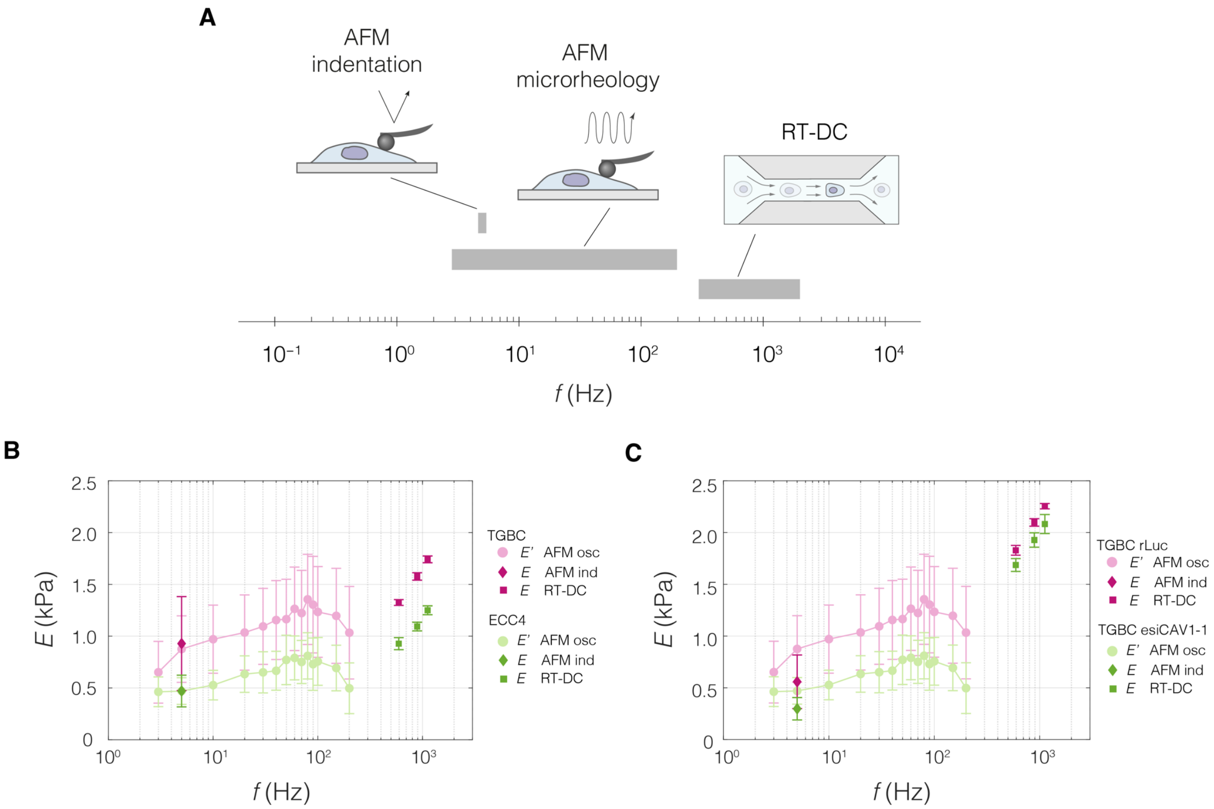


#### Figure 5–figure supplement 2

**Apparent Young's modulus values across the probing frequencies characteristic for the three measurement methods.**

(**A**) Graphical representation of measurement frequencies (the inverse of the time within which strain is induced) in the three methods for characterizing mechanical properties used in this study. For RT-DC, the frequency at which deformation is induced was deduced based on the time it takes to pass a 300-µm long square channel with an average velocity based on a range of flow rates typically used for 20 and 30-µm channels (see also **Table 6** and **Supplementary file 1**). (**B–C**) Apparent Young's moduli derived from RT-DC as well as AFM indentation and microrheology measurements plotted against probing frequency for ECC4 and TGBC cell lines (B), and CAV1 knock-down in TGBC cells (C). For RT-DC, data points for measurements at 0.16, 0.24 and 0.32 µl $s^{-1}$ flowrates are included for which the estimated frequencies, based on the time that it takes for the cell to pass through the channel, are equal to 593, 889, and 1119 Hz, respectively). For AFM microrheology, storage Young's moduli $E'$ were obtained from storage shear moduli ($G'$) according to the following equation: $E' = 2(1+\nu)G'$, assuming a Poisson's ratio, $\nu$, of 0.5. Datapoints correspond to means ± SD of individual cells (AFM microrheology) or medians ± MAD (of individual cells or measurement replicates in the case of AFM and RT-DC, respectively). Data corresponds to **Figure 5B-D** (B) and **Figure 5F-H** (C).

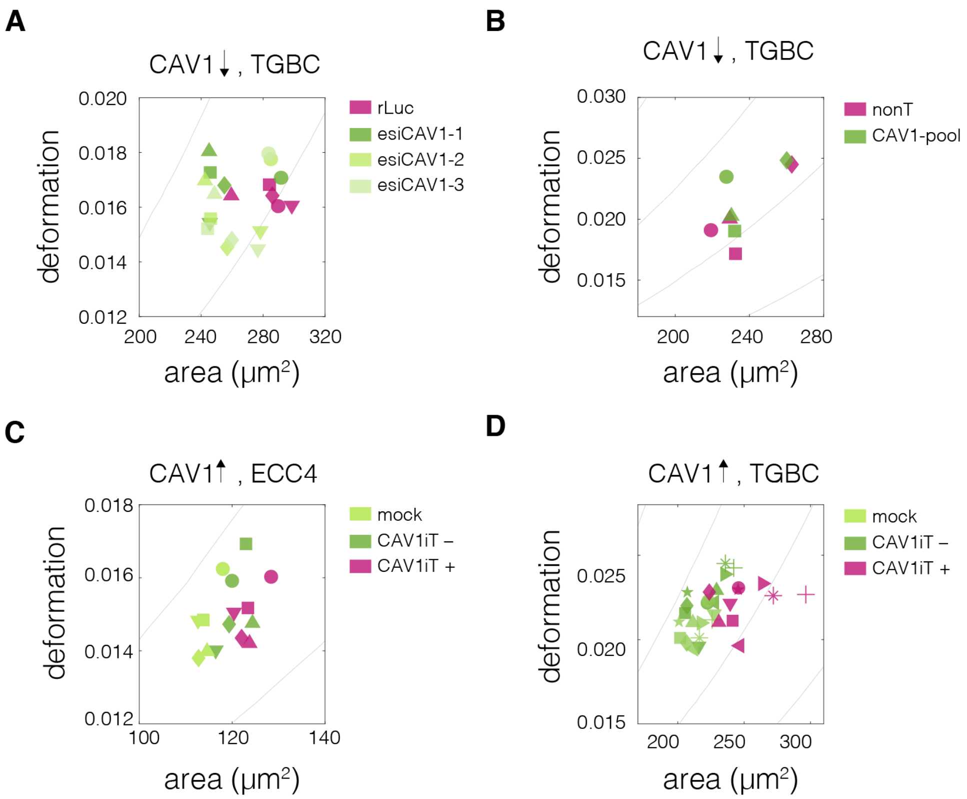


#### Figure 5–figure supplement 3

**Plots of area vs deformation from RT-DC measurements of cells with perturbed CAV1 levels.**

Panels correspond to the following experiments: (**A** and **B**) CAV1 knock-down in TGBC cells using esiRNA (A) and ONTarget siRNA (B), (**C** and **D**) transient CAV1 overexpression in ECC4 cells (C) and TGBC cells (D). Datapoints indicate medians of individual measurement replicates. The isoelasticity lines in the background (gray) indicate regions of same apparent Young's moduli. The symbol shapes represent experimental replicates.
