## Supplementary material for "De novo identification of universal cell mechanics gene signatures": Key Resources Table

| Reagent type (species) or resource | Designation | Source or reference | Identifiers | Additional information |
|---|---|---|---|---|
| Gene (*Homo sapiens, Mus musculus*) | *CAV1* | NA | HGNC:1527; MGI:102709 | caveolin 1 |
| Gene (*H. sapiens, M. musculus*) | *FHL2* | NA | HGNC:3703; MGI:1338762 | four and a half LIM domains 2 |
| Gene (*H. sapiens, M. musculus*) | *IGFBP7* | NA | HGNC:5476; MGI:1352480 | insulin like growth factor binding protein 7 |
| Gene (*H. sapiens, M. musculus*) | *TAGLN* | NA | HGNC:11553; MGI:106012 | transgelin |
| Gene (*H. sapiens, M. musculus*) | *THBS1* | NA | HGNC:11785; MGI:98737 | thrombospondin 1 |
| antibody | anti-Caveolin-1 (rabbit monoclonal) | Cell Signaling Technolo gy | CST: 3267; RRID: AB_2275 453 | WB (1:1000) |
| antibody | anti-GAPDH (rabbit polyclonal) | Abcam | Abcam: ab948 5; RRID: AB_3072 75 | WB (1:5000) |
| antibody | anti-rabbit HRP-conjugated (goat polyclonal) | Abcam | Abcam: ab970 69; RRID: AB_1067 9812 | WB (1:4000) |

| | | | | |
|---|---|---|---|---|
| cell line (*H. sapiens*) | Glioblastoma | *Poser et al., 2019* | X01; X04; X08 | human brain tumor cell lines; maintained in A. Androutsellis-Theotokis Lab (TU Dresden, Germany) |
| cell line (*H. sapiens*) | ECC4 | RIKEN BRC Cell Bank | RCB: RCB0982; RRID: CVCL_1190 | Intestine small-cell carcinoma; passage 7; medium: RPMI1640 (#11875093), 10% FBS |
| cell line (*H. sapiens*) | TGBC (TGBC18TKB) | RIKEN BRC Cell Bank | RCB: RCB1169; RRID: CVCL_3338 | intestine adenocarcinoma; passage 5; medium: DMEM (#11885084), 5% FBS |
| cell line (*H. sapiens*) | WA-hT | RIKEN BRC Cell Bank | RCB: RCB2279; RRID: CVCL_8766 | lung small-cell carcinoma; passage 54; medium: MEM (#11095080), 10% FBS |
| cell line (*H. sapiens*) | A549 | RIKEN BRC Cell Bank | RCB: RCB0098; RRID: CVCL_0023 | lung adenocarcinoma; passage 92; medium: DMEM (#11885084), 10% FBS |
| cell line (*H. sapiens*) | ECC10 | RIKEN BRC Cell Bank | RCB: RCB0983; RRID: CVCL_1188 | stomach small-cell carcinoma; passage 8; medium: RPMI1640 (#11875093), 10% FBS |
| cell line (*H. sapiens*) | MKN45 | RIKEN BRC Cell Bank | RCB: RCB1001; RRID: CVCL_0434 | stomach adenocarcinoma; passage 6; medium: RPMI1640 |

| | | | | |
|---|---|---|---|---|
| | | | | (#11875093), 10% FBS |
| cell line (*H. sapiens*) | MKN1 | RIKEN BRC Cell Bank | RCB: RCB1003; RRID: CVCL_1415 | stomach adenocarcinoma; passage 6; medium: RPMI1640 (#11875093), 10% FBS |
| cell line (*H. sapiens*) | MCF10A H1024R; MCF10A WT | *Juvin et al., 2013* | MCF10A H1024R; MCF10A WT | breast epithelial cells bearing single-allele oncogenic mutation of PIK3CA (H1024R); WT – isogenic control; kindly provided by L.R. Stephens (Babraham Institute, UK) |
| cell line (*H. sapiens*) | MCF10A-ER-Src | *Hirsch et al., 2009* | MCF10A ER-Src; RRID: CVCL_N 805 | breast epithelial cell model of TAM-inducible cancerous transformation driven by v-Src, a kind gift from K. Struhl (Harvard Medical School, MA, USA) |
| cell line (*M. musculus*) | iPSCs (F- and C-class) | *Urbanska et al., 2017* | iPSCs (F- and C-class) | induced pluripotent stem cells derived through reprogramming of murine fetal neural progenitor cells |
| cell line (*M. musculus*) | MEFs CAV1KO; MEFs WT | *Razani et al., 2001* | MEFs CAV1KO; MEFs WT | mouse embryonic fibroblasts derived from WT or CAV1KO littermate C57BL/9 mice; cell lines were a kind gift from M.P. Lisanti (University of |

| | | | | |
|---|---|---|---|---|
| | | | | Salford, Manchester, UK) |
| biological sample (*M. musculus*) | developing neurons (primary cells) | *Aprea et al., 2013* | developing neurons: PP – proliferating progenitors; NNs – newborn neurons | freshly isolated from double-reporter mouse line *Btg2*$^{RFP}$/*Tubb3*$^{GFP}$ by M. Dori in the Lab of F. Calegari |
| transfected construct (*H. sapiens*) | rLuc (esiRNA to rLuc) | Eupheria Biotech | Eupheria Biotech: RLUC | 200 ng per 2 µl RNAiMax in 12wp format |
| transfected construct (*H. sapiens*) | esiCAV1-1 (esiRNA to human CAV1, design 1, commercially available) | Eupheria Biotech | Eupheria Biotech: HU-03125-1 | 200 ng per 2 µl RNAiMax in 12wp format; see *Appendix X* for sequence details |
| transfected construct (*H. sapiens*) | esiCAV1-2 (esiRNA to human CAV1, design 2, custom) | Eupheria Biotech | Eupheria Biotech: HU-03125-2 | 200 ng per 2 µl RNAiMax in 12wp format; see *Appendix X* for sequence details |
| transfected construct (*H. sapiens*) | esiCAV1-3 (esiRNA to human CAV1, design 3, custom) | Eupheria Biotech | Eupheria Biotech: HU-03125-3 | 200 ng per 2 µl RNAiMax in 12wp format; see *Appendix X* for sequence details |
| transfected construct (*H. sapiens*) | nonT (ON-TARGETplus Non-targeting siRNA Pool) | Dharmacon | Dharmacon: D-001810-10-05 | 300 ng per 2 µl RNAiMax in 12wp format |
| transfected construct (*H. sapiens*) | CAV1-pool (ON-TARGETplus Human CAV1 siRNA, SMARTPool) | Dharmacon | Dharmacon: L-003467-00-0005 | 300 ng per 2 µl RNAiMax in 12wp format |

| | | | | |
|---|---|---|---|---|
| transfected construct (*H. sapiens*) | pCGIT-hCAV1 (plasmid, plasmid product referred to as CAV1iT) | this paper | pCGIT-hCAV1 | see *Plasmid for CAV1 overexpression* in *Materials and Methods*; plasmid map available on figshare (https://doi.org/10.6084/m9.figshare.28082240) |
| Chemical compound, drug | RNAiMax reagent | Thermo Fisher Scientific | Thermo Fisher Scientific: 13778030 | for siRNA transfections |
| Chemical compound, drug | Effectene transfection reagent | Qiagen | Qiagen: 301425 | for plasmid transfections |
| Chemical compound, drug | Methylcellulose | Alpha Aesar | Cat#: 036718.22 CAS 9004-67-5 | for preparation of viscosity-adjusted RT-DC measurement buffer |
| Software, algorithm | ShapeOut (version 1.0.10) | Zellmechanik Dresden | ShapeOut 1.0.10 | for analysis of RT-DC data, available on github: https://github.com/ZELLMECHANIK-DRESDEN/ShapeOut |
| Software, algorithm | JPK data processing software | JPK Instruments/ Bruker | *JPK DP* | for analysis of AFM experiments |
| Software, algorithm | PC-Corr network analysis | *Ciucci et al., 2017* | PC-Corr | code available on github: https://github.com/biomedical-cybernetics/PC-corr_net |

| | | | | |
|---|---|---|---|---|
| Software, algorithm | Cytoscape (version 3.8.0) | *Shannon et al., 2003* | cytoscape (RRID:SCR_003032) | available at https://cytoscape.org/ |
| Software, algorithm | Joint View Trustworthiness (JVT) | this paper | JVT | code available on github: https://github.com/biomedical-cybernetics/Joint-View-trustworthiness-JVT and figshare: https://doi.org/10.6084/m9.figshare.20123159 |
| Software, algorithm | Fiji, ImageJ | *Schindelin et al., 2012* | Fiji (RRID:SCR_002285) | https://fiji.sc/ |
| Other | PNP-TR-TL | Nanoworld, Switzerland | Nanoworld: PNP-TR-TL | tip-less AFM cantilevers, nominal spring constant $k = 0.08$ N m$^{-1}$ |
| Other | Arrow TL1 | Nanoworld, Switzerland | Nanoworld: Arrow TL1 | tip-less AFM cantilevers, nominal spring constant $k = 0.035$–$0.045$ N m$^{-1}$ |
| Other | polystyrene beads, 5-µm diameter | microParticles, Germany | microParticles: PS-R-5.0 | for decorating of the AFM cantilevers |
