## Supplementary File 1 for "De novo identification of universal cell mechanics gene signatures"

### Operation parameters of the three methods used for characterizing the mechanical properties of cells.

The table below summarizes the main parameters characteristic for the three methods applied for characterizing the mechanical properties of cells in this study: AFM indentation, AFM microrheology, and RT-DC. For AFM indentation, the setpoint of 2 nN leads to a resulting deformation on the order of 1 µm; for AFM microrheology, the deformation corresponds to the magnitude of the cantilever oscillations which was set to 10 nm. The corresponding strains are 7% for AFM indentation and 0.07% for AFM microrheology. For RTDC, a mean absolute strain on the order of 17% was reported (*Urbanska et al., 2020*), this corresponds to ca 2.5 µm deformation. For the above estimation, the cell diameter was assumed to be 15 µm (ECC4 and TGBC cells have a cross-section area within100–300 µm$^2$ (see **Figure S2B**), corresponding to 11.3–19.5 µm diameter). For RT-DC, the probing frequency was deduced based on the time it takes to pass a 300-µm long square channel with an average velocity based on a total flow rate of 0.16 µl s$^{-1}$ and 30 µm channel width (a setting most frequently used in this study, see Table S5), For AFM microrheology, the frequency is equivalent to the oscillation frequency; and for AFM indentation, it is deduced from the extension speed of 5 µm s$^{-1}$ and an average deformation of 1 µm.

| | **AFM indentation** | **AFM microrheology** | **RT-DC** |
|---|---|---|---|
| **cell state** | adherent | adherent | suspended |
| **probing lengthscale** | local (5-µm bead) | local (5-µm bead) | whole-cell |
| **frequency (probing timescale)** | 5 Hz (200 ms) | 3 Hz (300 ms) – 200 Hz (5 ms) | ~ 600 Hz (1.67 ms) |
| **induced deformation** | 1 µm | 10 nm | 2.5 µm |
| **strain** | 7 % | 0.07 % | 17 % |
| **deformation rate** | 5 µm s$^{-1}$ | 0.03–2 µm s$^{-1}$ | 1.5 mm s$^{-1}$ |
| **strain rate** | 0.35 Hz | 0.002–0.140 Hz | 102 Hz |
