## Supplementary File 4 for "De novo identification of universal cell mechanics gene signatures"

### Joint-view trustworthiness (JVT) pseudocode and computational complexity analysis

#### *Pseudo-code of JVT*

$DS = \{ds_i, ds_2, \ldots, ds_n\}$, – set of names of the datasets to load. Each $ds_i$ is composed of 4 variables: the corresponding $x_i$, $common\_gene\_names_i$, $sample\_lables_i$, $AUC\_label_i$.

$x_i$ – the actual dataset within each $ds_i$, where features/genes are on rows and samples are on columns. Each $x_i$ is organized having the same genes and in the same order on the rows

$common\_gene\_names_i$ – an array of gene names common to each dataset

$sample\_lables_i$ – a 2D array where each row corresponds to $x_i$ in each $ds_i$ reporting for each sample the name of one of the two groups to which it belongs in $x_i$

$AUC\_label_i$ – an array containing the name of one of the two groups of samples that should be used as positive class for AUC-ROC evaluation; each $AUC\_label_i$ corresponds to each $ds_i$

$CoMa$ – an array containing the names of genes used for computing the combinatorial marker

$al$ – a numerical array of the same length as $CoMa$ containing one of the values in $\{1, -1\}$ for each gene, indicating whether alignment is needed $(-1)$ or not $(1)$

$norm$ – type of gene normalization involved. Offered options: '*log10*' and '*z-score*'

$T$ – an integer indicating the number of times the samples are drawn to build null model distributions

$Z$ – length of $common\_gene\_names$

$C$ – length of $CoMa$

$S$ – size of $DS$

$N_i$ – number of samples of a $ds_i$

$result$ – $2 \times (C + 1)$ matrix, where rows 1 and 2 store minimum AUC score and p-value of minimum AUC score, respectively, as computed by JVT. Each column corresponds to each marker (single+combinatory).

---

**Alg1. Joint Multiview Trustworthiness** $JVT$ $\quad O(S(ZN + (T + C)(CZ + CN)) + C(Z + T + S + C) + TS)$

---

1. **Input** = datasets $DS = \{ds_i, ds_2, \ldots, ds_n\}$, a Combinatorial Marker array $CoMa$, an array $al$, normalization type $norm$, sampling times $T$
2. find gene indices, $ind\_CoMa$, in $CoMa$ via any $ds_i$ using its $x_i$ and $common\_gene\_names_i$ // $O(ZC)$

   // *initialisation of variables begins*
3. generate a series $l = 1\ to\ Z$ // *used as indices to datasets* ; $O(Z)$
4. $te \leftarrow set\_difference(l, ind\_CoMa)$ // $te$ *contains* $l$ *with* $ind\_CoMa$ *removed* ; $O(ZC)$
5. $f \leftarrow length(te)$
6. append $ind\_CoMa$ back to $l$ // $O(C)$

**if** $T < (Z - C)$ **OR** $T > (Z - C)$ **then** *// for randomisation; if $T = Z - C$, no need to randomise*

$rand \leftarrow$ uniformly generate $T$ random integers $\in [1, f]$ *// $O(T)$*

$l \leftarrow te(rand)$ *// $l$ is updated and now contains random indices from $te$ and has length $= T$*

append $ind\_CoMa$ back to $l$ *// $O(C)$*

**end if**

$L = length(l)$ *// $L = T + C$*

initialise $L \times S$ zero matrix, $ROC\_area$, and $(T + 1) \times S$ zero matrix, $ROC\_area\_comb$

*// $ROC\_area$ and $ROC\_area\_comb$ are used to store single and combinatory gene AUC performance,*
*// respectively, and have the same number of columns as the no. of datasets*

*// $O(S(T + C))$ for $ROC\_area$ matrix and $O(S(T + 1))$ for $ROC\_area\_comb$ matrix*

*// initialisation of variables ends*

**for** $k = 1$ **to** $S$ **do** *// for each dataset* ; $O(S(ZN + (T + C)(CZ + CN))$

load dataset $ds = DS(k)$ *// this loads all the variables, $x$, $common\_gene\_names$,*
*// $sample\_lables$ and $AUC\_label$ corresponding to dataset, $ds$*

perform *log* or *z-score* normalisation to $x$ corresponding to $ds$ based on $norm$
*// if $norm$ does not contain offered options, use log normalisation as default;* $O(ZN)$

**for** $i = 1$ *to* $L$ **do** *// for all sampling times $T$ and $C$*

*// $O((T + C)(ZC + CN + C + 4N)) \approx O((T + C)(CZ + CN))$*

pick the $i^{th}$ index in $l$ and select the corresponding gene, $rand\_gene$ in $x$

$AUC = AUC\_ROC(rand\_gene, sample\_lables, AUC\_label)$

*// compute $AUC\_ROC$ of $rand\_gene$ in $x$ using the corresponding*
*// $sample\_lables$ and $AUC\_label$* ; $O(N)$

$ROC\_area(i, k) = max(AUC, 1 - AUC)$ *// for single gene* ; $i$ = *row*, $k$ = *column*

**if** $i \leq (T + 1)$ **then** *// for combinatory gene*

**if** $i \leq (T)$ **then**

$rand =$ uniformly generate $C$ random integers $\in [1, f]$ *// $O(C)$*

$R = te(rand)$

*// $R$ contains random indices from $te$ and has length $= C$*

**elseif** $i = (T + 1)$ **then**

$R = ind\_CoMa$ *// no randomisation; only marker indices used*

**end if**

$xt \leftarrow$ genes identified by $R$ indices in $x$ *// $O(ZC)$*

**for** $sd = 1$ **to** $C$ **do** *// for gene alignment ;* $O(C(N + N)) \approx O(CN)$

**if** $al(sd) == -1$ **then**

$m = mean(xt(sd))$ // $O(N)$

// *take the mean of the gene identified by its* $\mathbf{sd}$ *index*

$t(sd)) = 2 \times m - xt(sd))$ // *align and update gene* // $O(N)$

**end if**

**end for**

$xt = column\_mean(xt)$ //*take the column-wise mean of* $xt$ *genes* ; $O(CN)$

$AUC = AUC\_ROC(xt, sample\_lables, AUC\_label)$

// *compute* $AUC$ *of* $xt$ *using the corresponding* $\mathbf{sample\_lables}$ *and*
// $AUC\_label$ ; $O(N)$

$ROC\_area\_comb(i, k) = max(AUC, 1 - AUC)$

**end if**

**end for**

**end for**

$ROC\_area\_min$ = compute the minimum of $ROC\_area$ along each row // $O(S(T + C))$

// $ROC\_area\_min$ is a vector containing the minimum AUC performance of single gene
// across the datasets

$ROC\_area\_comb\_min$ = compute minimum of $ROC\_area\_comb$ along each row // $O(S(T + 1))$

// $ROC\_area\_min$ is a vector containing the minimum AUC performance of combinatorial
// gene across the datasets

initialise $2 \times (C + 1)$ zero matrix, $result$ // *for result compilation* ; $O(2(C + 1))$

**for** $i = 1$ **to** $C$ **do** // *for single marker* ; $O(C(Z + T + C))$

$result(1, i) = ROC\_area\_min$ value corresponding to the index $ind\_CoMa(i)$ // $O(Z)$

$result(2, i) =$ (number of times $ROC\_area\_min \geq result(1, i))/L$ // *p-value* ; $O(T + C)$

**end for**

$result(1, C + 1) = ROC\_area\_min\_comb(T + 1)$ // *for combinatory marker*

$result(2, C + 1) =$ (number of times $ROC\_area\_min \geq result(1, C + 1))/(T + 1)$

// *p-value* ; $O(T + 1)$

append *'Comb Marker'* to $CoMa$

set row names of $result$ = {'min AUC-ROC', 'JVT p-value'}

set column names of $result = CoMa$

**Output** = $result$

*Computational complexity of JVT*

We begin with the breakdown of computational complexity of JVT as presented in *Alg1* and then provide the overall associated complexity of JVT. From line 2 in *Alg1*, we can see the complexity of finding $C$ number of marker genes in a set of $Z$ number of common genes is $O(ZC)$. Similarly, lines 3, 4 and 6 have the complexity of $O(Z)$, $O(ZC)$ and $O(C)$, respectively. Line 8, where we generate $T$ random numbers and has the complexity of $O(T)$, where $T$ represents the sampling times, and line 9 runs at $O(C)$. In line no. 13, we create two zero matrices for storing the results, $ROC\_area$ and $ROC\_area\_comb$, having the computational complexity of $O(S(T+C))$ and $O(S(T+1))$, respectively, where $S$ is the number of datasets involved. The loop in line 14 has a total complexity of $O(S(ZN+(T+C)(CZ+CN))$. This complexity can be arrived using lines 15 through 40, which compose the statements of the loop, as follows. Line no. 16, which normalizes each value in a given dataset has a complexity of $O(ZN)$. Line 17, which is also a loop, has a total complexity of $O((T+C)(CZ+CN))$. This can be arrived at by inspecting lines 19, 23, 28, 31, 32, 35 and 36. While, the complexities of these lines are easy to deduce and are given in the algorithm itself, lines 19, 29 and 36 are of particular interest here. Since line 19 computes the $AUC-ROC$ score of the gene with respect to the number of samples the gene carries, $N$, it has the computational complexity of $O(N)$. The same can be said of line 36. Line 19 is a loop and performs the gene alignment with a complexity of $O(CN)$. Putting all this together, we get the computational complexity of line 14 stated above. Furthermore, line 41, which computes the minimum $ROC\_area$ of all the sampled genes, $T+C$, across all the datasets, $S$, has the complexity of $O(S(T+C))$. Similarly, for line 42, we have $O(S(T+1))$. Line 43 creates a $2\times(C+1)$ zero matrix and intuitively runs at $O(2(C+1))$, and loop in line 44 at $O(C(Z+T+C))$. Finally, line 49 has the computational complexity of $O(T+1)$. Summing up the algorithm's complexity and simplifying, we get,

$$O(S(ZN+(T+C)(CZ+CN))+C(Z+T+S+C)+TS).$$

Clearly, if the values $S$, $N$ and $C$ are highly dominated by the $T$ and $Z$, which is the case in our study, the effect of $S$, $N$ and $C$ in the overall complexity of the JVT becomes negligible. Hence, the complexity boils down to $O(Z+T+ZT) \approx O(ZT)$, i.e., the JVT's complexity is non-linear in $Z$ and $T$.
