## Supplementary File 5 for "De novo identification of universal cell mechanics gene signatures"

### Sequences of esiRNAs used for CAV1 knock-down experiments

>HU-03125-1 (414 bp)

GAGCTGAGCGAGAAGCAAGTGTACGACGCGCACACCAAGGAGATCGACCT
GGTCAACCGCGACCCTAAACACCTCAACGATGACGTGGTCAAGATTGACT
TTGAAGATGTGATTGCAGAACCAGAAGGGACACACAGTTTTGACGGCATT
TGGAAGGCCAGCTTCACCACCTTCACTGTGACGAAATACTGGTTTTACCG
CTTGCTGTCTGCCCTCTTTGGCATCCCGATGGCACTCATCTGGGGCATTT
ACTTCGCCATTCTCTCTTTCCTGCACATCTGGGCAGTTGTACCATGCATT
AAGAGCTTCCTGATTGAGATTCAGTGCATCAGCCGTGTCTATTCCATCTA
CGTCCACACCGTCTGTGACCCACTCTTTGAAGCTGTTGGGAAAATATTCA
GCAATGTCCGCATC

>HU-03125-2 (325 bp)

CCAAAATGTTGGTCATTTTATGTTAAGGGAAGAATTCCAGGGTATGGCCA
TGGAGTGTACAAGTATGTGGGCAGATTTTCAGCAAACTCTTTTCCCACTG
TTTAAGGAGTTAGTGGATTACTGCCATTCACTTCATAATCCAGTAGGATC
CAGTGATCCTTACAAGTTAGAAAACATAATCTTCTGCCTTCTCATGATCC
AACTAATGCCTTACTCTTCTTGAAATTTTAACCTATGATATTTTCTGTGC
CTGAATATTTGTTATGTAGATAACAAGACCTCAGTGCCTTCCTGTTTTTC
ACATTTTCCTTTTCAAATAGGGTCT

>HU-03125-3 (680 bp)

GAGTTGCTGCAAACCTGACCCCTGCTCAGTAAAGCACTTGCAACCGTCTG
TTATGCTGTGACACATGGCCCCTCCCCCTGCCAGGAGCTTTGGACCTAAT
CCAAGCATCCCTTTGCCCAGAAAGAAGATGGGGGAGGAGGCAGTAATAAA
AAGATTGAAGTATTTTGCTGGAATAAGTTCAAATTCTTCTGAACTCAAAC
TGAGGAATTTCACCTGTAAACCTGAGTCGTACAGAAAGCTGCCTGGTATA
TCCAAAAGCTTTTTATTCCTCCTGCTCATATTGTGATTCTGCCTTTGGGG
ACTTTTCTTAAACCTTCAGTTATGATTTTTTTTTCATACACTTATTGGAA
CTCTGCTTGATTTTTGCCTCTTCCAGTCTTCCTGACACTTTAATTACCAA
CCTGTTACCTACTTTGACTTTTTGCATTTAAAACAGACACTGGCATGGAT
ATAGTTTTACTTTTAAACTGTGTACATAACTGAAAATGTGCTATACTGCA
TACTTTTTAAATGTAAAGATATTTTTATCTTTATATGAAGAAAATCACTT
AGGAAATGGCTTTGTGATTCAATCTGTAAACTGTGTATTCCAAGACATGT
CTGTTCTACATAGATGCTTAGTCCCTCATGCAAATCAATTACTGGTCCAA
AAGATTGCTGAAATTTTATATGCTTACTGA
